# Transcriptomic atlas of premalignant oral squamous cell carcinoma in an aging mouse model reveals an enhanced immune response and dysregulation of head and neck tissue stem cells

**DOI:** 10.64898/2026.03.20.713236

**Authors:** Heidi Kletzien, Nguyen-Ahn Nguyen, Siddhartha G. Jena, Jason D. Buenrostro, Amy J. Wagers

## Abstract

Oral squamous cell carcinomas (OSCC) account for ∼90% of all oral malignancies and have devastating effects on overall health and quality of life. However, little is known about the early initiating events that drive the development of oral leukoplakia-like premalignant lesions (OPLs) and disease progression. Here, we create a mouse model of tobacco-related premalignant OSCC that takes into consideration its primary risk factors, including advancing age and male sex. This model notably recapitulates the age variant patterns of OSCC risk observed in humans, with a higher prevalence of oral premalignant lesions in older mice. In addition, by building a transcriptomic atlas with this system, we reveal genetic signatures associated with oncogenic progression in the tongue and buccal epithelium, and their resident somatic tissue stem cells. We also identify several novel transcriptomic signatures of premalignancy in OSCC, including enhanced immune response and expansion and dysregulation of head and neck tissue stem cells. These findings offer a new framework for investigating physiologically-relevant risk factors and drivers of OSCC and illuminate novel biological pathways underlying its pathology.

## Introduction

Cancers of the head and neck are a heterogenous group of malignancies affecting epithelial, mucosal, and skeletal tissues of the upper aerodigestive tract. In the United States, head and neck cancer (HNC) causes more than 500,000 deaths annually,^1-3^ with primary risk factors that include advancing age,^4-6^ male sex,^7,8^ and exposure to environmental carcinogens, particularly tobacco.^9^ These risk factors are expected to drive increasing incidence of HNC over the next decade, with particular impact on oral squamous cell carcinoma (OSCC) - one of the most prevalent HNCs.^10,11^

Management of OSCC typically involves multimodal treatment strategies that are relatively ineffective, suffer high rates of tumor recurrence, and can have devastating effects on speech, swallowing, and overall quality of life.^6,12-15^ The significant burden that OSCC places on individuals, and on the healthcare system more broadly,^16,17^ emphasizes the urgent need for new, more effective treatment approaches, which in turn will depend on expanding our currently limited knowledge of the critical molecular circuits and biological mechanisms that initiate and propel these highly aggressive malignancies.

The etiologic role of tobacco use in HNC, including OSCC, is clear and has been well studied. Tobacco contains more than 70 known carcinogens and 5,000 different chemicals with carcinogenic activity.^18^ Among these, the nitrosamines, NNK and NNN (4-[methlnitrosamino]-1-[3-pyridyl]-1-butanone and N’-nitrosonornicotine, respectively) are the most carcinogenic.^18^ Both NNK and NNN are tobacco-specific cytotoxic and genotoxic agents. Exposure to these agents generates DNA adducts that cause single-strand breaks (SSB), insertions and deletions (INDELs), and base-pair substitutions.^19,20^ Carcinogen-induced SSBs are particularly mutagenic and, unless repaired by intrinsic cellular processes, can lead to the activation of oncogenes (i.e., *H-ras, K-ras*), inactivation of tumor suppressors (i.e., *p53)*, and tumor induction.^21-23^ Both NNK and NNN exposures correlate strongly with OSCC risk and diagnosis.^24-27^

Thus far, mechanistic studies of OSCC have focused predominantly on the genomic, proteomic, and metabolic properties of late-stage tumors and patient-derived HNC cell lines, with relatively few illuminating the early underlying mechanisms that drive oncogenic transformation.^23,28-33^ This situation likely reflects the difficulty that exists in obtaining premalignant lesions and tissue biopsies from human subjects at precancerous timepoints.^34^ In preclinical animal studies, one of the most utilized and robust models of tobacco-related OSCCs involves exposure of mice to the chemical carcinogen, 4NQO (4-Nitroquinoline N-oxide), which generates reactive oxygen species and DNA adducts that cause mutagenesis and drive tumor induction, similar to NNK and NNN.^35-37^ Typically, the 4NQO model of OSCC involves administration of 4NQO to young mice in their drinking water, although 4NQO may alternatively be painted on to the tissue(s) of interest. In either case, administration of carcinogen is typically continuous over a 16-week treatment period, with animals monitored for subsequent tumor development for an additional 12 weeks. The 4NQO model reproducibly generates invasive squamous cell carcinomas of the tongue and esophagus, presenting typically at around 28-weeks after initial 4NQO exposure. The majority of published studies to date have focused on these late-stage tumors, generally in young male mice.^38-41^

Recognizing the strength of this OSCC model, we sought to leverage it to investigate the dynamics of early, initiating events in OSCC evolution, while also considering additional known human epidemiological variables, including the impact of advancing age and genetic sex, on tumor induction. In the current study, we track tumor development following exposure to 4NQO across age and sex, developing a comprehensive cell biological and transcriptomic atlas that profiles early genetic alterations and changes in tissue-resident stem cell populations that are involved in OSCC emergence in the tongue and buccal epithelium of young, middle-aged, and aged male and female mice. These data offer a detailed understanding of the dynamics of early initiating events in OSCC evolution and implicate a number of novel targets and pathways in the malignant transformation of cells in these tissues. We anticipate that this atlas will serve as a resource for the field, illuminating particularly the early events in cellular transformation and tumor emergence and aiding in the development of precision-based interventions that target the underlying biology of OSCC in its premalignant state.

## Results

### 4NQO-induced oral premalignant lesions emerge at an accelerated rate in aged animals and mimic initiation of human oral squamous cell carcinoma

To investigate the effects of increasing age and genetic sex on the development of oral leukoplakia (white patch-like oral premalignant lesions [OPLs] of the tongue that are the most common precursor of OSCC), young (4 months old), middle-aged (18 months old), and aged (24 months old) C57BL/6J male and female mice were administered the tobacco mimetic 4NQO (4-Nitroquinoline N-oxide) in their drinking water for 2-, 4-, or 8-weeks (Fig. 1A). Mice were examined weekly for the development of lesions on the ventral and dorsal surfaces of the tongue and for lesions of the buccal epithelium, through to the study endpoint. No lesions of the buccal epithelium were observed. Kaplan-Meier lesion-free survival analysis (log rank, Mantel-Cox) was conducted to compare the main effects of age or sex, alone, on the development of tongue OPLs. Lesion-free survival differed significantly among the three age groups, χ^2^(2)=8.407, p=0.015, with the shortest mean time to lesion development observed in the aged cohort (6.62 weeks, 95%CI, 6.1 -7.1) and longest for the young group (7.69 weeks, 95%CI, 7.7–8) (Fig. 1B). The middle-aged group exhibited an intermediate time to lesion development (7.26 weeks, 95%CI, 6.8 -7.7). A log rank test to discriminate sex-specific differences in the lesion-free survival distribution did not indicate statistically significant differences (χ^2^(1)=2.243, p=0.134); however, the data do suggest a trend for accelerated lesion development in males (7.06 weeks; 95%CI, 6.7-7.4) compared to females (7.27 weeks; 95%CI, 6.9-7.7) (Fig. 1B). A Cox proportion hazards regression was also performed to assess potential interaction effects between age and sex on lesion development. The -2 Loglikelihood value of 686.384 and the omnibus test of model coefficients indicate a good model fit (χ^2^(2)=5.414, p=0.067). Hazard ratios indicate that increasing age is a significant predictor of lesion development (B=0.278, p=0.045, HR=1.32, 95%CI [1.0-1.7]), while male sex is not (B=-0.218, p=0.352, HR=0.8, 95%CI [0.5-1.3]).

**Fig. 1.**
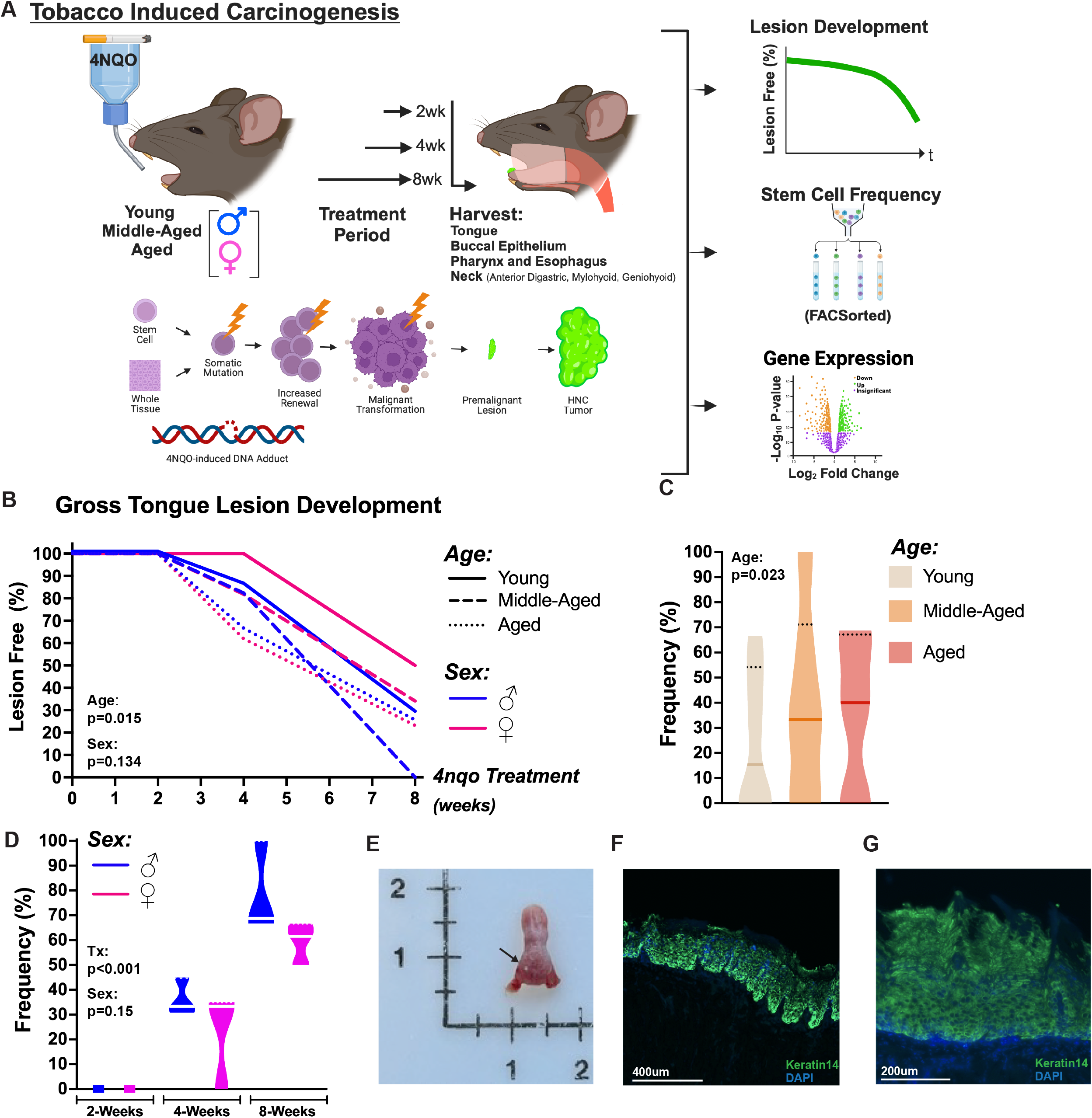
(A) Schema for Experimental Design. Young, middle-aged, and aged male and female mice were randomized into no treatment (regular water control condition) or 2-week, 4-week, or 8-week 4NQO treatment periods. Animals were monitored for lesion development. Histologic, Cytofluorometric, and/or Transcriptomic analyses were performed immediately following study completion. (B) Kaplan-Meier lesion free-survival curve comparing the main effects of age or sex, alone, on the development of tongue OPLs. Binomial logistical regression show that age (C) and treatment period (D) are significant predictor variables of OPL development. (E) Representative photo-micrograph of OPL (arrow) on dorsal surface of tongue in a 4NQO-treated C57BL/6 mouse. Representative immunofluorescence images of an OPL from a 4NQO-treated mouse show thickening of the tongue epithelium (Keratin 14).

We next performed a binomial logistic regression to ascertain the effects of age, sex, and 4NQO treatment period on the likelihood that mice present with OPLs at the study endpoint. The logistic regression was statistically significant, (χ^2^(3)=84.43, p<0.001). The model explains 45.1% (Nagelkerke R^2^) of the variance and accurately classifies 75.9% of OPLs in 4NQO-treated mice. Sensitivity was 65.5%, while specificity was 81.6%. The positive and negative predictive values were 66.7% and 81.0%, respectively. Of the predictor variables, only age (p=0.023; 95%CI, 1.1–2.5) (Fig. 1C) and treatment period (p<0.001; 95%CI, 4.4-13.9) (Fig. 1D) were statistically significant. Aged mice had a 1.65 times higher odds ratio to have OPLs at study endpoint, and longer 4NQO treatment time (in weeks) was associated with a 7.8 times increased likelihood of OPL development. Although not significant, female sex was associated with a lower chance of developing OPLs (p=0.15; 95%CI, 0.3-1.2; HR=0.6) compared to mice of male sex. To assess the regression model’s performance, we also applied ROC (receiver operating characteristic) curve analysis. The binary model exhibited excellent discrimination of mice with OPLs and mice without lesions, with an area under the curve of 0.847 (95%CI, 0.8-0.9) (Supp. 1). Taken together, this comprehensive data set suggests that the 4NQO-induced premalignant mouse model of OSCC exhibits a disease-relevant partial sex-bias and appropriately recapitulates the age-variant patterns observed in human patients, with a significantly higher prevalence of oral premalignant lesions in aged mice (Fig. 1).

### Transcriptomic atlas of premalignant head and neck tissues and their resident tissue stem cells in aged, 4NQO-exposed, male and female mice

The etiologic role of tobacco use in late-stage human OSCCs is well understood, as are the genomic and transcriptomic changes of late-stage 4NQO-induced OSCCs in, primarily, young male mice. Studies of such late-stage tumors have identified activated oncogenes, inactivated tumor suppressors, and dysregulation of pathways involved in cell proliferation, differentiation, apoptosis, and immunosuppression in OSCC tumors, with several pathways commonly affected in humans and model organisms. However, mechanistic studies of OSCC malignancy thus far have focused predominantly on the properties of well-established tumors, cell lines, and patient-derived primary cells, and offer relatively little information regarding the early, premalignant changes that occur across lifespan and with potentially divergent manifestations on the basis of genetic sex. Likewise, there has been little exploration to date of the specific involvement of resident head and neck tissue stem cells in the development of OSCC following carcinogenic insult, and it remains uncertain whether oral tissue-resident stem cells can serve as tumor cells-of-origin for OSCC initiation.

To begin to fill these gaps in existing knowledge, we utilized the 4NQO model to create an atlas detailing the early, premalignant changes in transcriptomic states of head and neck tissues and their resident somatic tissue stem cell populations in young, middle-aged, and aged male and female mice, with and without 4NQO treatment. We chose an 8-week timepoint for 4NQO exposure based on our observation that OPLs were observed in an average of 66.6% of mice after this duration of 4NQO exposure (Fig. 1).

We performed bulk RNA-seq and differential gene expression analysis of the whole tongue and whole buccal epithelium, and of FACSorted epithelial stem and progenitor cells, mesenchymal stromal stem and progenitor cells, and satellite cells (muscle stem cells) isolated from tissues of carcinogen-exposed and control mice. Stem cell populations were identified and isolated using well-validated cell surface marker profiles.^42-47^ Following normalization and removal of low-abundance transcripts, principal component analysis (PCA) of differentially expressed genes revealed that gene expression clustered predominately by tissue and stem cell population type (Fig. 2A, E), and by treatment condition (Fig. 2B, Supp. 2A, D, G, J, M). However, some overlap was detected in gene expression profiles when looking at age (Fig. 2C, Supp. 2B, E, H, K, N) and sex (Fig. 2D, Supp. 2C, F, I, L, O; Supp. 3) while also controlling for treatment condition (8-weeks 4NQO treatment versus no treatment). This expression similarity across different age groups and between genetic sexes may indicate that 4NQO exposure is the primary driver of the variation in gene expression observed in our atlas of OSCC premalignancy. For these reasons, we decided to analyze global transcriptomic changes first between the 8-week 4NQO and no treatment conditions across all tissue types and stem cell populations.

**Fig.2.**
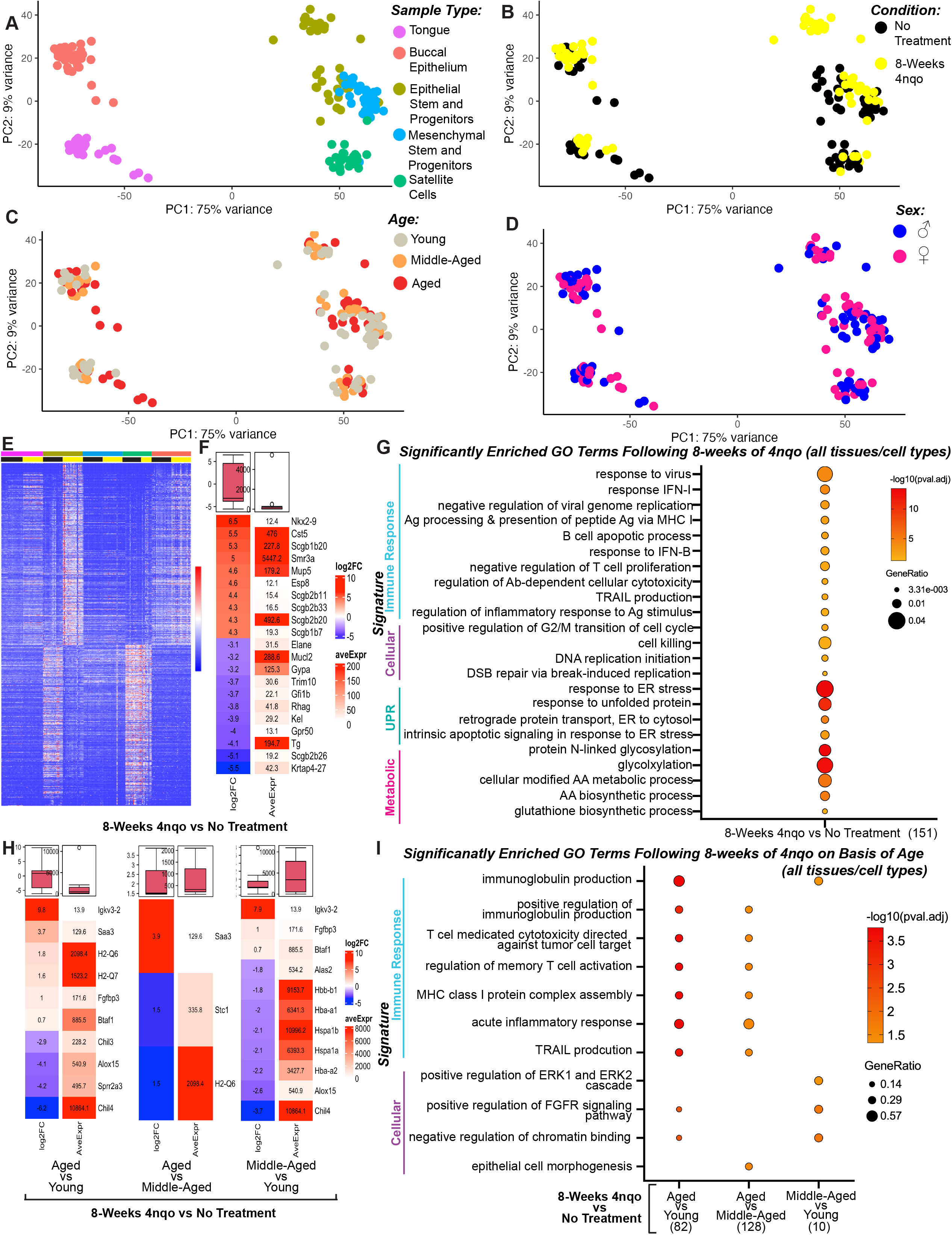
Principal component analysis (PCA) plots of bulk RNA-sequencing show that gene expression profiles cluster primarily by tissue and stem cell populations types (A) and by treatment condition (B) and not by age (C) and/or sex (D). (E) k-Means clustering of transcripts differentially expressed in the 4NQO treatment metagroup compared to the no treatment control metagroup. (F) Top 10 most significantly upregulated and downregulated genes in the 4NQO metagroup compared to the no treatment control metagroup. (G) Significantly enriched gene ontology terms in the 4NQO treatment metagroup compared to the no treatment control metagroup. (H) Top significantly upregulated and downregulated genes based on 4NQO treatment condition and age. (I) Significantly enriched gene ontology terms in the 4NQO treatment metagroup compared to the no treatment control metagroup and based on age.

Using DESeq2, we identified 1,366 significantly up-regulated genes and 1,318 significantly down-regulated genes in the 4NQO treatment metagroup compared to the no treatment control metagroup (padj<0.05) (Fig. 2E,F). Focusing on the top ten most significantly up-regulated genes in this particular data set, the secretoglobin (SCGBs, padj<0.0046, FDR<0.05) gene family showed abundant representation (Fig. 2F). While little is known about the role of secretoglobins in OSCC, abnormal expression profiles of SCBGs have been observed in other cancer types, including breast, lung, and colorectal cancers, and SCBGs have been suggested as potential biomarkers and prognostic indicators in cancers of epithelial origin.^48-56^ Cst5, a candidate tumor suppressor in colon cancers, also exhibited elevated expression within this data set (log_2_FC=5.48, padj=1.2363E-08, FDR<0.05), which may point to an early and globally protective role of this protein through its function inhibiting proteases involved in cancer cell invasion and metastasis (Fig. 2F).^57-59^ SMR3A (log_2_FC=4.96, padj=1.7155-08, FDR<0.05), a gene that is highly expressed and may serve as a potential prognostic marker in head and neck squamous cell carcinomas (HNSCCs), oropharyngeal squamous cell carcinomas (OPSCCs), and adenoid cystic carcinomas of the head and neck, exhibited the highest average expression across all tissues and cell types in early 4NQO-associated OSCC premalignancy.^60-63^ Gene ontology (GO) analysis evaluating all differentially expressed genes (padj<0.05, q-value =1) revealed enrichment of 350 biological processes. To ameliorate potential overlapping gene sets and terms, we also performed an additional clustering algorithm, which reduced the list to 151 significantly enriched GO terms (Fig 2G). Among the top GO terms, four themes emerged that can be summarized into the following enrichment groups: immune response, cellular processes, unfolded protein response, and metabolic processes (Fig. 2G). All of these biological processes are highly relevant in cancer and shared among the tissue and cell types profiled in this atlas.

To investigate global transcriptomic changes across the different tissue types and stem cell populations that might be influenced by age and 4NQO exposure in our study, we employed a design formula in DESeq2 that allowed examination of the treatment condition (8-weeks 4NQO treatment versus no treatment) based on age and its interaction, controlling for our specific age comparisons of interest (aged versus young, aged versus middle-aged, and middle-aged versus young). When comparing age groups of interest, we identified nine significantly up-regulated genes in aged versus young mice exposed for 8-weeks to 4NQO, as compared to untreated control mice. We also identified three significantly up-regulated genes in aged versus middle-aged and middle-aged versus young comparisons (Fig. 2H). Of the most significantly expressed genes (padj<0.05), Igkv3-2 (ortholog human IGKV3-20),^64,65^ H2-Q7,^65-68^ and Saa3^69^ appear to be conserved across several age comparisons, with the most common overlap occurring between the aged versus young and middle-aged versus young comparisons. These data suggest a heightened immune response and proinflammatory state that associates with increasing age in early stages of OSCC development (Fig. 2H), which is also reflected in observed meta-group GO signatures, appearing first at the middle-aged versus young comparison, and persists across both aged comparisons (Fig. 2H, 2I). After correcting for redundant GO terms and gene sets, enriched GO terms clustered primarily into two thematic groups involved again with enhanced immune response and upregulation of cell-based processes (Fig. 2I). These data reveal discrete changes in cell states occurring early in the tumorigenic process and suggest an important influence of tissue remodeling that is consistent with lesion development and tumor progression.

### Immune signature predominates transcriptomic profiles in aged 4NQO-treated mice in premalignancy

Along with tobacco use, advancing age is a primary risk factor for the development of OSCC. However, the genes and pathways contributing to OSCC development as a function of age remain largely unexplored. To investigate transcriptomic profiles that may contribute to the accelerated development of oral premalignant lesions in aged and middle-aged animals, compared to young (Fig. 1), we employed the same design formula in DESeq2 mentioned above, but focused instead on individual tissue types and stem cell populations in isolation. GO analysis of each tissue and stem cell population revealed a striking inflammatory response that was enhanced in aged versus young and aged versus middle-aged comparisons. In some comparisons, immune response-related categories were the only GO terms that were significantly enriched or accounted for more than 90% of enriched terms (e.g., aged versus young epithelial stem and progenitors) (Fig. 3A). The immune-related terms that predominated in these analyses were involved in antigen processing and presentation pathways, MHC protein complexes, T and B cell mediated processes, and enhanced cytokine and immunoglobulin production. Immune-related gene and protein families that were highly expressed in aged tissues and cell types included MHC-Is (H2-Q6, H2-Q7, H2-K1, H2-T10, H2-M9, H2-DMb1,) and cytokines (IFNβ, TNF, Il1a, Il1r2, Ccl12, Ccl1, Ccl28, Cxcl10, Ccr2).^65,70-75^ When comparing enriched immune-related GO terms across tissue and stem cell types by age, separately, the tongue and buccal epithelium showed the most significantly enhanced immune signature across all age comparisons, followed by the significant enrichment of immune GO signatures in epithelial stem and progenitor cells and in mesenchymal stromal stem and progenitor cells in the aged versus young and aged versus middle comparisons (Fig. 3A). Interestingly, the only significant immune GO signature for muscle stem cells (also known as satellite cells) was observed in the aged versus young comparison (Fig 3A). These data reveal an enhanced and heterogenous age-related immune response across tissues and cell types.

**Fig. 3.**
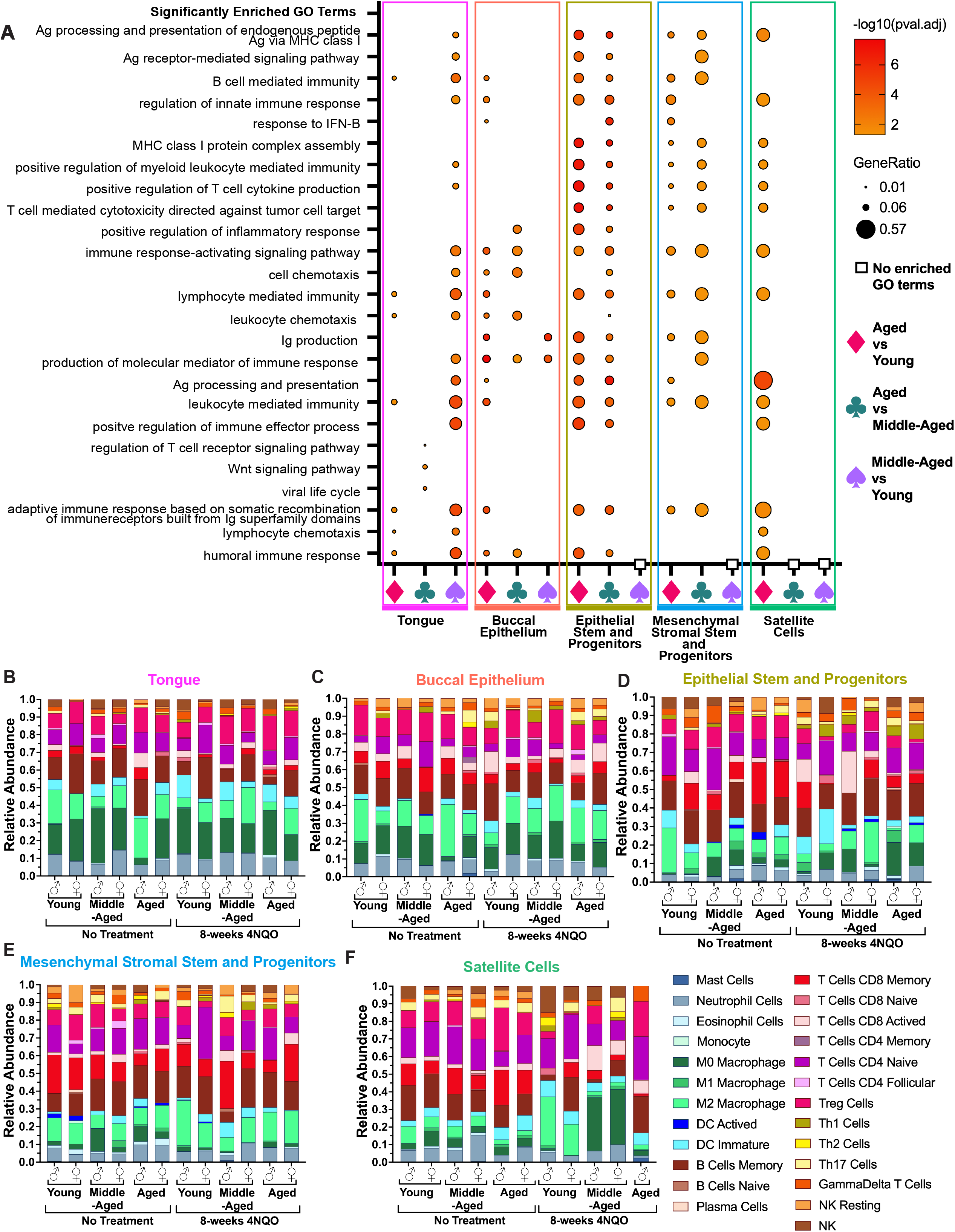
(A) Significantly enriched gene ontology terms in the 4NQO treatment group compared to the no treatment control group and based on age and tissue and cell type. Digital cytometry and deconvolution of bulk sequenced tissues and stem cell populations using CIBERSORTx of the tongue (B), buccal epithelium (C), epithelial stem and progenitor cells (D), mesenchymal stem and progenitor cells, and satellite cells (E).

Given the critical role that immune cells play in cancer initiation, we also examined immune cell and immunologic marker composition specifically by performing digital cytometry and deconvolution on our bulk sequenced tissues and stem cell populations using CIBERSORTx (Fig. 3B-F).^76^ For the tongue, relative abundance profiles for all immune cell types showed significant main effects for lymphocytes (Age, p=0.026), T cells (Age, p=0.029), and dendritic cells (Sex, p=0.044); significant interaction effects for CD8 T cells (8-weeks 4NQO treatment by Sex, p=0.019; 8-weeks 4NQO treatment by Age by Sex, p=0.077), activated CD8 T cells (8-weeks 4NQO treatment by Age by Sex, p=0.025), naïve CD8 T cells (8-weeks 4NQO treatment by Sex, p=0.005), memory CD8 T cells (8-weeks 4NQO treatment by Sex, p=0.045), and naïve CD4 cells (8-weeks 4NQO treatment by Age by Sex, p=0.048) (Fig. 3B). A trend towards significance was observed for myeloid cells (8-weeks 4NQO treatment by Sex, p=0.061; Age by Sex, p=0.077), macrophages (8-weeks 4NQO treatment, p=0.061), B cells (8-weeks 4NQO treatment by Sex, p=0.082), T regulatory cells (8-weeks 4NQO treatment, p=0.054), and T helper 17 cells (8-weeks 4NQO treatment, p =0.056)(Fig. 3B). For the buccal epithelium, relative abundance profiles showed significant interaction effects for macrophages (8-weeks 4NQO by treatment Sex, p=0.027) and T helper cells (8-weeks 4NQO treatment by Sex, p=0.025), and a trend towards significance for lymphocytes (8-weeks 4NQO treatment by Sex, p=0.062) and T cells (8-weeks 4NQO treatment by Sex, p=0.062) (Fig. 3C). Epithelial stem and progenitor cells, mesenchymal stem and progenitor cells (Fig. 3 D-E), and satellite cells (Fig. 3F) also displayed immune cell marker signatures. These data highlight a common induction of immune responses upon tissue damage, likely linked to 4NQO exposure and the initiation of cellular transformation, which may aid in the recruitment of immune cells.^77^

### 4NQO-induced alterations in head and neck tissue stem cell frequency and transcriptomic profiles in aging male and female mice

Genotoxic events in resident tissue stem cells play a critical role in the development of many cancers. In head and neck tissues, aging-associated and carcinogen-induced DNA strand breaks can result in early transcriptomic changes that trigger the expansion and/or dysregulation of resident stem cells. However, the specific carcinogenic effects of 4NQO on head and neck tissue stem cells, and how these intersect with aging in OSCC pathogenesis, is as yet unknown. To uncover potential changes in the proliferative dynamics of tissue stem cells following 4NQO treatment, epithelial and mesenchymal stromal stem and progenitor cell, and satellite cell frequencies were analyzed from the oral epithelium (Fig. 4A-B, Supp. 4A), tongue (Fig. 4C-F), neck muscles (Supp. 4B-C), and the pharynx and esophagus (Supp. 4D-E) after 4NQO treatment periods of 2-, 4-, or 8-weeks.^42-45,47,78^ The frequencies of each stem cell population were then compared to the frequencies of analogous populations in unexposed control conditions. Three-way ANOVAs were performed to assess the interaction effects of the 4NQO treatment condition (4NQO treatment [2-, 4-, or 8-weeks] or no treatment control), age (young, middle-aged, or aged), and genetic sex (male or female) on stem cell frequency. To account for multiple comparisons, the Sidak correction was applied. For epithelial stem and progenitor cells (EpSCs) isolated from the oral epithelium, there was a statistically significant three-way interaction between treatment condition, age, and sex (F(6, 95)=2.948, p=0.027) (Fig. 4A-B). Simple-simple comparisons revealed significantly higher frequencies of EpSCs in all age groups in both males and females for all 4NQO treatment conditions, with peak stem cell frequency occurring after 2-weeks or 4-weeks of 4NQO treatment. At the 8-week time point, EpSC frequency remained elevated compared to control mice, but was reduced relative to the 2- and 4-week 4NQO treatment timepoints. These early changes in EpSC frequency may indicate enhanced proliferation and possible clonal expansion within this stem cell population in response to 4NQO exposure. Compared to young female mice, young male mice had a lower frequency of EpSCs at the 4-week (p=0.032) timepoint and a higher frequency at 8-weeks (p=0.026). A significant main effect for age was observed in mesenchymal stromal stem and progenitor cells (F(6, 104)=4.889, p=0.009) in the absence of any other significant main effects or interaction effects (Fig. 4C-D). Young mice had higher frequencies of mesenchymal stromal stem and progenitor cells compared to middle-aged (p=0.011) and aged (p=0.072) mice.

**Fig. 4.**
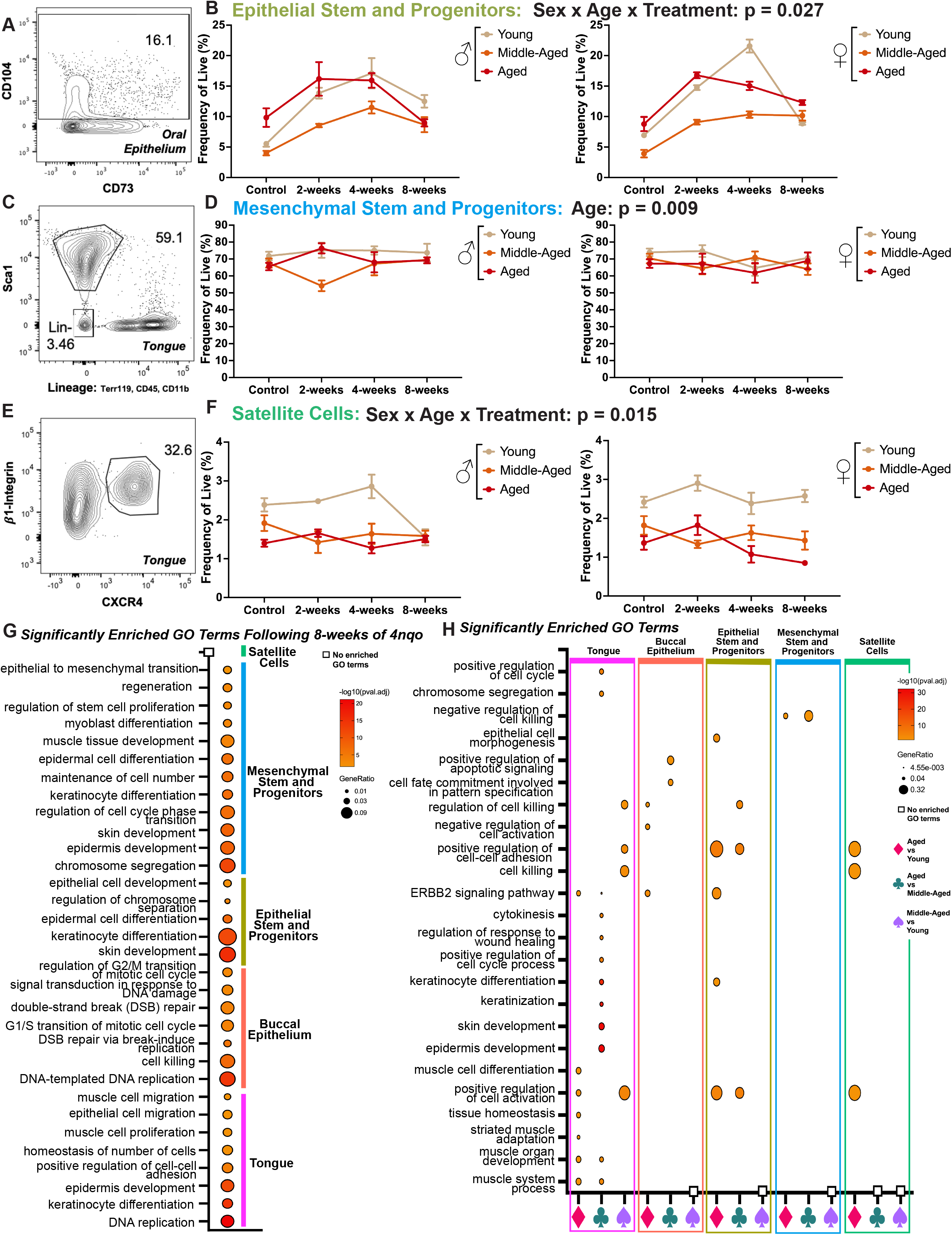
Representative flow cytometry plots and frequency of live summary plots for epithelial stem and progenitor cells (A, B), mesenchymal stem and progenitor cells (C, D), and satellite cells (E, F) (G) Gene ontology analysis of significantly enriched terms in the 4NQO treatment group compared to the no treatment control group by tissue and cell type. (H) Significantly enriched gene ontology terms in the 4NQO treatment group compared to the no treatment control group and based on age by tissue and cell type.

A significant three-way interaction effect was also observed for satellite cells (SCs) isolated from the tongue F(2, 104)=2.783, p=0.015) (Fig. 4E-F). Simple-simple comparisons showed statistically significant alterations across age groups and for each sex at all 4NQO exposure timepoints. Young mice overall had a higher SC frequency compared to all other age groups across all experimental conditions, while in young mice, peak SC frequency occurred at the 4-week 4NQO timepoint for males, and at 2-weeks for females. In middle-aged male and female mice, peak SC frequency occurred at the 4-week 4NQO timepoint, and in aged male and female mice, SCs peaked at the 2-week 4NQO treatment timepoint. Compared to young female mice, young male mice had a lower frequency of SCs (p=0<0.001), whereas aged males had a higher SC frequency (p=0.031) at the 8-week timepoint compared to females.

We also observed significant interaction effects in epithelial stem and progenitor cells (age by treatment: F(6,95)=5.492, p<0.001), in satellite cells isolated from neck muscles (age by treatment: F(6,108)=4.201, p<0.001), in pooled pharyngeal and esophageal satellite cells (age by treatment: F(6,106)=3.278, p=0.005); we observed main effects in neck mesenchymal stromal stem and progenitor cells (MSCPs) (age: F(2,108)=9.68, p<0.001) and in pooled pharyngeal and esophageal MSCPs (treatment: F(3,107)=15.938, p<0.001) (Supp. 4). Taken together, these data suggest that even following a very brief exposure to the tobacco mimetic 4NQO, ranging from only 2-to 8-weeks, alterations arise in multiple tissue-resident stem cell populations found in head and neck tissues. Most striking is the 4NQO-induced expansion of these stem cell populations across age groups, which differed in male versus female mice. This observation suggests that damage induced by 4NQO is intrinsic to either the stem cells themselves or to their local milieu, which contributes indirectly to an alteration in stem cell frequency as compared to age- and sex-matched controls.

Using DESeq2, we next investigated potential cellular and molecular regulators that may have contributed to the observed alterations in stem cell frequency induced by 4NQO exposure. GO analysis revealed the enrichment of many biological processes relevant to stem cell proliferation, expansion, and differentiation following 8-weeks of 4NQO exposure (Fig. 4G). Of note, shared GO processes across tissue types and stem cell populations included the enrichment of pathways related to epithelial development and expansion (e.g. epidermis and skin development; epithelial cell and keratinocyte development, differentiation, and migration), tissue remodeling (e.g., regeneration, muscle development, stem cell proliferation, maintenance of cell number, cell killing, and epithelial to mesenchymal transition), and cell-based mechanisms that may be contributing to observed changes (e.g., chromosome segregation, replication, state transitions in the cell cycle, DNA damage, and double strand break repair).

We additionally employed a design formula in DESeq2 to examine biological processes in the context of age following 8-weeks of 4NQO exposure (Fig. 4I). Significant enrichment of GO terms related to stem cell expansion and tissue remodeling were most apparent in the aged versus young comparison, and included shared pathways related to positive regulation of cell activation, positive regulation of cell-cell adhesion, keratinocyte differentiation, and upregulation of the ERBB2 signaling pathways (Fig 4I). Interestingly, the enrichment of gene pathways related to apoptosis and cell killing were only observed in the tongue and buccal epithelium, which may indicate a global response to the dysregulation and expansion of resident stem cell populations in these tissues following 4NQO-mediated damage. There were no significantly enriched GO terms in the middle aged-versus young comparisons for the buccal epithelium, epithelial stem and progenitor cells, or mesenchymal stromal stem and progenitor cells related to cell expansion or changes in cell/tissue architecture. This observation may suggest that aged (versus young and versus middle aged) head and neck tissues and their resident stem cell populations are the most sensitive to 4NQO-induced damage and implicate these cells as an underlying cause for the observed acceleration of oral premalignant lesions in this model of OSCC.

## Discussion

In this study, we established a mouse model of premalignant tobacco-related oral squamous cell carcinoma (OSCC) that convincingly reflects the age- and sex-related emergence pattern of OSCC initiation in human patients, with oral premalignant lesions (OPLs) occurring more frequently in older mice, and a trend towards accelerated OPL development in males. We further generated a comprehensive transcriptomic atlas from this model that profiled early gene expression alterations occurring in the tongue and buccal epithelium and their resident somatic tissue stem cells. Our comprehensive analysis revealed that gene expression programs cluster predominately by tissue and stem cell population type and by the 4NQO (4-Nitroquinoline N-oxide) exposure condition, with the emergence of four thematic gene signatures related to enhanced immune response, upregulated cellular processes that regulate different tissue/cell states, changes in metabolic states, and activation of the unfolded protein response. This study fills an important gap in knowledge by revealing the dynamic progression of transcriptional changes underlying early initiating events in OSCC development and implicating several specific targets, pathways, and transcriptomic signatures in the malignant transformation of stem cell populations within these tissues.

In mice, the 4NQO model is the most well-established and well-characterized mouse model for understanding the *in vivo* biology of tobacco-related OSCCs.^36,79^ One of the major advantages of the 4NQO chemical carcinogen model is its ability to mimic the stepwise progression of human OSCC from healthy normal tissue to the development of oral leukoplakic-like premalignant lesions and tumors that share histologic features associated with all stages of human disease including hyperplasia and dysplasia, papillary squamous tumors, carcinomas *in situ*, and invasive squamous cell carcinomas of the tongue.^35,36,80,81^ Even more striking are the similarities between the genomic signatures and mutational landscapes of high-grade and late-stage 4NQO induced dysplasias and tumors of the dorsal tongue and advanced human OSCCs from smokers. Indeed, recent studies report signature associations of between 90 to 94% among these mouse and human tumor samples.^39,79^ Although overwhelmingly complex and heterogenous, both mouse and human tumor analyses have shown overlapping tobacco-related signatures in late-stage tumors, more specifically the activation of oncogenes, mutations in tumor suppressor genes, and the dysregulation of pathways involved in cell proliferation, differentiation, and apoptosis, immunosuppression, and the upregulation of many proinflammatory pathways. However, more recently there has been a focus on deciphering the early mutagenic and inflammatory responses of OSCC initiation to better understand disease progression through the study of oral leukoplakic-like premalignant lesions and the surrounding tissue environment.^39,41,82-85^

Oral premalignant lesions (OPLs) are precursors to OSCCs and strongly associate with tobacco use.^82^ Approximately 30% of OPLs progress to invasive carcinomas and yet, few studies to date have focused on this early stage of tumor development, which represents an intermediary phase of OSCC initiation and progression.^84,86-90^ Results from our mouse model of premalignant OSCC, the first to incorporate age and sex as biological variables in the 4NQO mouse model, recapitulated OPL emergence patterns observed in humans. This feature further demonstrates the value of this mouse model in mapping the evolution of early initiating events in OSCC. To better understand mechanisms contributing to the emergence of OPLs, we focused our analysis on global transcriptomic changes occurring in the whole tongue and buccal epithelium and on signatures that might be driving the malignant transformation and dysregulation of resident stem cell populations, rather than on the OPLs themselves.

After 8-weeks of 4NQO exposure, we identified several highly expressed, premalignant transcriptomic markers in the 4NQO treatment metagroup that have previously been associated with cancer initiation, including the secretoglobin (SCGB) gene family,^48-56^ Nkx2-9 (ortholog to human NKX2-8),^91-95^ and SMR3A ^62^,^60,61,63^ as well as 6 genes (Gpr50, Gfi1b, Trim10, Gypa, Mucl2 [ortholog to human MUC2], Elane) that are known to be downregulated in colorectal, lung, breast, laryngeal, and head and neck squamous cell carcinomas. Interestingly, when we take age into consideration in analyzing the 4NQO treatment metagroup, we observe conserved pro-inflammatory (Igkv3-2, H2-Q genes, Saa3)^65,96^ and tumorigenic (up: Fgfbp3, Btaf1; down: Alox15, Hba and Hbb gene families, Chil3/4)^97-107^ signatures across multiple age group comparisons. Collectively, these dysregulated genes may serve as potential biomarkers for OSCC initiation and warrant further investigation due to their known involvement in other cancer types. However, unlike other studies in human and mouse that profiled OPLs, exclusively, we did not observe overtly significant changes in expression of Trp53, Notch1, Cdkn2a, Fat1, Pcna, Egfr, or Mmps in our tissues and cell types of interest. This lack of overlap in expression profiles may highlight the distinct alterations that occur in premalignancy to drive subsequent pro-oncogenic changes that contribute to OPL development and progression to OSCC. Indeed, our data suggest that early alterations in the immune and stromal environment may facilitate OPL development, and proposes a novel role for epithelial, mesenchymal stromal, and muscle stem and progenitor cells in these dysregulated tissues, potentially priming healthy cells for malignant transformation.

We also observed an expansion of epithelial stem and progenitor cells that had immune-related expression profiles following 4NQO exposure. The abnormal proliferation and expansion of epithelial cells, in particular, has been linked to tumor promoting processes, including the infiltration of macrophages and myeloid cells, and angiogenesis.^11,26^ Strikingly this 4NQO-induced, age-related signature and altered differentiation profile was not limited to the epithelial stem and progenitor populations alone, as we also observed an enhanced and striking immunophenotypic signature in the mesenchymal stromal stem and progenitor cells, and in satellite cells of aged mice. Taken together, these global changes could contribute to immune evasion at this stage of disease progression and to the accelerated development of OPLs in these aged animals compared to their young counterparts.

In summary, we developed a comprehensive transcriptomic atlas that profiles early genetic alterations involved in OSCC emergence in young, middle-aged, and aged male and female mice in the tongue and buccal epithelium, and in their resident somatic tissue stem cells. This premalignancy model strikingly mimics human OSCC development and provides a detailed understanding of the dynamics of early initiating events in OSCC evolution, highlighting several novel targets and pathways involved in the malignant transformation of cells in these tissues. These results open a unique window into the early transcriptomic changes and cellular genetic events that contribute to tumor development, revealing crucial biological underpinnings of OSCC in its premalignant state.

## Methods

### Mice

All experimental procedures were approved and performed in compliance with Harvard University’s Institutional Animal Care and Use Committee (IACUC, Protocol Number 11-26). Male and female mice were maintained in a pathogen-specific free facility accredited by the Association and Accreditation of Laboratory Animal Care (AALAC) and housed in standard ventilated racks at a density of 2-5 mice per cage, provided food and water *ad libitum*, and kept on a 12-hour light/dark cycle. C57BL/6 mice were either purchased from Jackson Laboratory (JAX, #000664) and/or bred, or obtained from the National Institute on Aging Aged Rodent Colony (Charles River Laboratories). Mice that were bred from those originally purchased from JAX, were maintained in-house for a minimum of 8 weeks prior to experimentation with littermate controls. All NIA obtained mice were acclimated to the mouse facility for a minimum of 2 weeks prior to experimentation.

### Oral Squamous Cell Carcinoma Induction

Young (2-4 mo. old; n=94), middle-aged (16-18 mo. old; n=89) and aged (22-24 mo. old n=116) male (n=167) and female (n=144) C57BL/6J mice were randomized into no treatment (regular water control condition, n=89) or 4NQO (4-Nitroquinoline N-oxide, Millipore Sigma; 4nqo treated water) treatment (n=212) conditions (Fig 1A). Mice randomized into the 4NQO treatment conditions, were administered 4NQO in drinking water, diluted to 100 μg/mL, for either 2-week (n=65), 4-week (n=72), or 8-week (n=75) treatment periods. 4NQO drinking water was prepared and changed every 1 to 2 weeks and stored in light-shielded water bottles. Once a week, mice were weighed and anesthetized by isoflurane (1-3%) inhalation to screen the oral cavity for lesions of the tongue and buccal epithelium.

Following the experimental duration, mice were euthanized and the head and neck tissues (tongue [pooled extrinsic and intrinsic muscles], oral and buccal epithelium, pharynx and esophagus (pooled), and neck [pooled digastric, mylohyoid and geniohyoid muscles]) were dissected for downstream analyses.

### Cell Isolation and Fluorescence Activated Cell Sorting (FACS)

Epithelial stem and progenitor cells, mesenchymal stromal progenitor cells, and satellite cells (muscle stem cells) were isolated and FACSorted as described previously. Briefly, the head and neck tissues (tongue [extrinsic and intrinsic muscles], oral and buccal epithelium, pharynx and esophagus, and neck [digastric, mylohyoid and geniohyoid muscles]) were enzymatically and mechanically digested with collagenase and dispase and further isolated via centrifugation. All cells were subjected to ACK Lysis on ice for 5 minutes. For epithelial stem cell and progenitor cell isolation, cells were stained with an antibody mix (FITC anti-CD104 [β4-Integrin, 1:25] or PE-Cy7 anti-CD104 [β4-Integrin, 1:50], PE anti-CD73 [1:50], APC-Cy7 anti-Terr119 [1:200], APC-Cy7 anti-CD11b [Mac1, 1:200], APC-Cy7 anti-CD45 [1:200]) on ice, in the dark for 45 minutes. Calcein blue was used to identify live cells. Live (Calcein blue^+^) Terr119^-^ CD45^-^ CD11b^-^ β4-Integrin^+^ CD73^+^ were FACSorted on a BD FACS ARIA II. For mesenchymal stromal progenitor cell and satellite cell isolation, cells were stained with an antibody mix (PE anti-CD29 [β1-Integrin, 1:100], Biotin anti-CD184 [CXCR4, 1:100], APC anti Sca1 [1:200], APC-Cy7 anti-Terr119 [1:200], APC-Cy7 anti-CD11b [Mac1, 1:200], APC-Cy7 anti-CD45 [1:200]) on ice, in the dark for 30 minutes, followed by secondary (Streptavidin-PE-Cy7 [1:200]) for 20 minutes. Live cells were identified with calcein blue. Sca1 expressing mesenchymal progenitors (Live Ter119^-^ CD45^-^ CD11b^-^ Sca1^+^) and muscle stem cells (Live Ter119^-^ CD45^-^ CD11b^-^ Sca1^-^ B1-Integrin^+^ CXCR4^+^) were then FACSorted on a BD FACS ARIA II. Cells were sorted into TRIzol LS Reagent (Invitrogen). Analysis of FACSorted stem cell populations was performed using FlowJo software.

### RNA-seq cell and tissue preparation and analysis

For FACSorted stem cell populations, RNA was isolated using TRIzol LS (Invitrogen) and RNeasy Micro columns (Qiagen) on a Qiacube. For whole tissues samples (tongue and buccal epiethelium), tissues were homogenized in TRIzol (Invitrongen) in gentleMACS C Tubes on a gentleMACS dissociator (Miltenyi Biotec) and then RNA isolated using RNeasy mini columns on a Qiacube. Total RNA quality and integrity was then assessed using RNA 6000 pico and nano kits, respectively, on a 2100 Bioanalyzer System (Agilent) or using high sensitivity RNA ScreenTape or RNA ScreenTape assays, respectively, on a 4200 TapeStation (Agilent). Smart-Seq v4 (Takara Bio) libraries were prepared, indexed (Nextera XT), and sequenced on an Illumina NovaSeq 6000 (2x S1 flow cells) with paired-end reads.

Base calls were converted to FASTQ format using bcl2fastq, and FASTQ reads were aligned to the mouse genome (GENCODE GRCm38/mm10) using STAR. Aligned reads were annotated to both exons and introns using featureCounts. DESeq2 v1.46.0 was used normalize raw read counts and remove low counts, and to quantify gene-level expression between conditions (control vs 8-weeks 4nqo treatment), age (aged vs young, aged vs middle-aged, and middle-aged vs young), and sex (male vs female). Multi-factor design formulas to model specific interactions effects were also incorporated to quantify gene expression changes on the basis of condition and age, and condition and sex. Principal component analysis was performed to visualize and assess variation between sample/tissue/cell type, condition, age, and sex. An adjusted p-value of 0.05 was used to account for multiple hypothesis testing. Genes with a FDR ≤0.05 for whole tissues and genes with an a FDR ≤0.1 for FACSorted stem cell populations were deemed significant. Gene Ontology analysis was performed using clusterProfiler on significantly differentially regulated genes (q-value=0.1, and FDR ≤0.05 for whole tissues and FDR ≤0.1 for FACSorted stem cell populations) for all comparisons noted above. Heatmaps were generated using Morpheus (Broad Institute) and the immune cell compositions for all samples were deconvoluted using CIBERSORTX and a reference immune cell matrix for mouse.

### Histology

For histological analyses, tongues were embedded in OCT, fresh-frozen in liquid nitrogen chilled 2-mehthylbutane, and longitudinally sectioned at 12 μm using a Leica Biosystems cryostat. Sagittal sections were either fixed in formalin and stained with Hematoxylin and Eosin by standard methods. Sections were imaged using an ECHO Confocal (4x or 20x).

### Statistical Analyses

Data are presented as means ± SE. IBM SPSS software was used to perform all statistical tests unless stated otherwise: Kaplan-Meier curve, COX regression, Binomial logistic regression, Receiver operating characteristic curve, or three-way ANOVA. Results with a p-value <0.05 were considered statistically significant. IBM SPSS, GraphPad Prism, and RStudio software programs were used to generate data plots.

## Supporting information

Merged Supplemental Files and Legends

## Data Availability

The datasets generated and/or analyzed during the current study will be made available in a public data repository (GEO) upon publication. All other data are available within the Article and Supplementary Data, or are available from the authors upon request.

## Acknowledgements

This work was supported in part by NIH grants DP1 AG048917 (to A.J.W.), F32 AG071208-01(to H.K.), and P30 AG031679 (Boston Claude D. Pepper Older Americans Independence Center Research Education Component Award, to H.K), as well as funding from the Harvard Stem Cell Institute, and the Paul F. Glenn Center for Biology of Aging Research at Harvard Medical School. S.G.J. and J.D.B. were funded by NIH awards R01AR083416 and R01AR080110. We thank J. LaVecchio, N. Kheradmand, and G. Kassis for flow cytometry assistance, the Bauer Core Facility for sequencing services, and Robert Shekoyan for animal husbandry services and help with histology.

## Author Contributions

Conceptualization, H.K., N.N., and A.J.W.; Methodology, H.K., N.N., and A.J.W.; Data Curation, H.K., S.G.J, J.D.B; Investigation, H.K., N.N.; Resources, A.J.W and J.D.B; Software, H.K. S.G.J, N.N; Writing, H.K., S.G.J, A.J.W; Funding Acquisition, H.K. J.D.B, A.J.W

