## Supplementary material for "Transcriptomic atlas of premalignant oral squamous cell carcinoma in an aging mouse model reveals an enhanced immune response and dysregulation of head and neck tissue stem cells": Merged Supplemental Files and Legends

**Supplemental Figures (Extended Data)**

**Supp. 1**

ROC (receiver operating characteristic) curve analysis to asses performance of the binomial regression model.

**Supp. 2**

Principal component analysis (PCA) plots of bulk RNA-sequencing for each tissue and stem cell type (tongue, A-C; buccal epithelium, D-F; epithelial stem and progenitors (G-I), mesenchymal stem and progenitor cells (J-L), and satellite cells (M-O) by treatment condition (A, D, G, J, M), age (B, E, H, K, N), and sex (C, F, I L, O).

**Supp. 3**

(A) k-Means clustering of transcripts differentially expressed in the 4NQO treatment group compared to the no treatment control group based on sex. (B) Top significantly upregulated and downregulated genes in the 4NQO group compared to the no treatment control group based on sex. (G) Significantly enriched gene ontology terms in the 4NQO treatment group compared to the no treatment control group on the basis of sex.

**Supp. 4**

Frequency of live summary plots for epithelial progenitor cells (A), mesenchymal stem and progenitor cells (B), and satellite cells (C) from pooled neck muscles, and mesenchymal stem and progenitor cells (D) and satellite cells (E) from pooled esophagus and pharynx.

Supp. 1

A

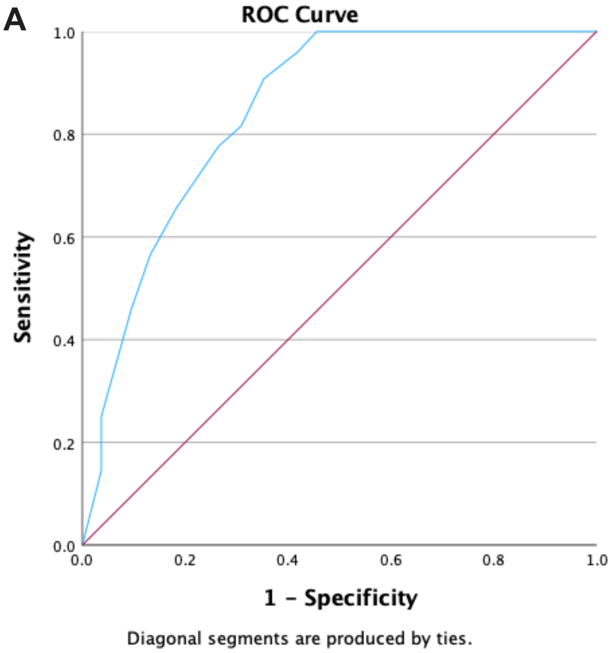
ROC Curve
Sensitivity
1.0
0.8
0.6
0.4
0.2
0.0
0.0
0.2
0.4
0.6
0.8
1.0
1 - Specificity
Diagonal segments are produced by ties.

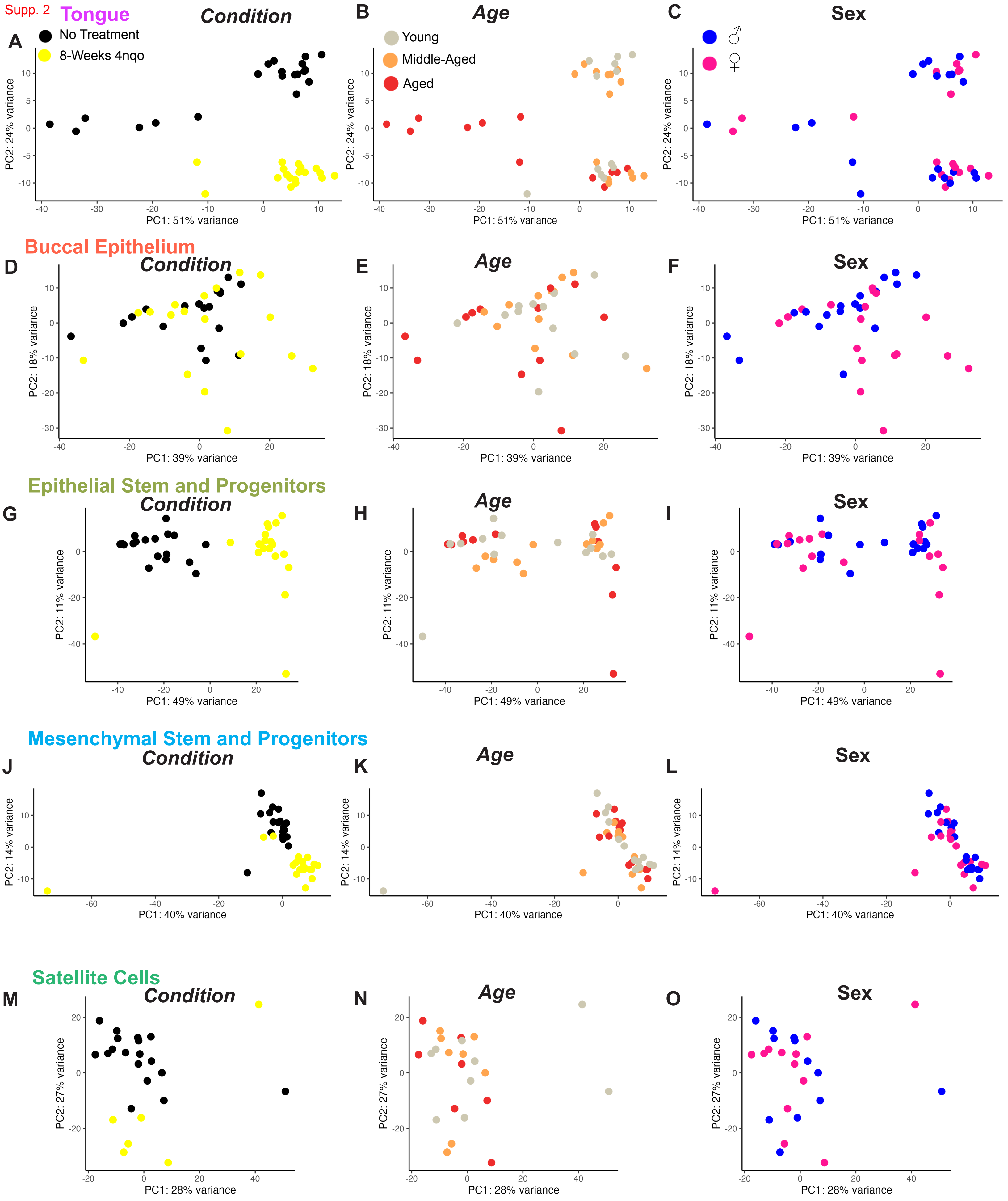
Supp. 2
Tongue
Condition
No Treatment
8-Weeks 4nqo
Age
Young
Middle-Aged
Aged
Sex
♂
♀
PC2: 24% variance
PC1: 51% variance
Buccal Epithelium
PC2: 18% variance
PC1: 39% variance
Epithelial Stem and Progenitors
PC2: 11% variance
PC1: 49% variance
Mesenchymal Stem and Progenitors
PC2: 14% variance
PC1: 40% variance
Satellite Cells
PC2: 27% variance
PC1: 28% variance
A
B
C
D
E
F
G
H
I
J
K
L
M
N
O

Supp. 3

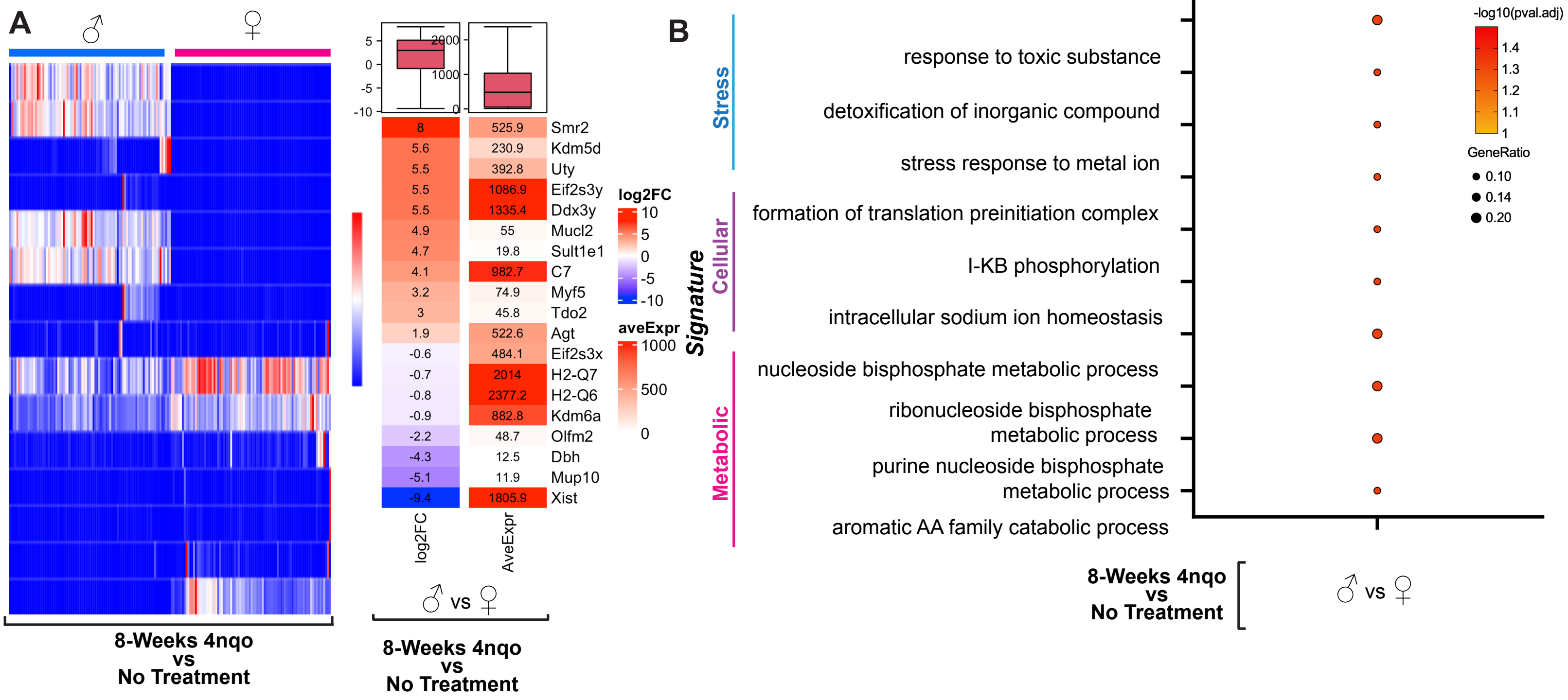

A
♂
♀
8-Weeks 4nqo
vs
No Treatment
log2FC
10
5
0
-5
-10
aveExpr
1000
500
0
Smr2 8 525.9
Kdm5d 5.6 230.9
Uty 5.5 392.8
Eif2s3y 5.5 1086.9
Ddx3y 5.5 1335.4
Mucl2 4.9 55
Sult1e1 4.7 19.8
C7 4.1 982.7
Myf5 3.2 74.9
Tdo2 3 45.8
Agt 1.9 522.6
Eif2s3x -0.6 484.1
H2-Q7 -0.7 2014
H2-Q6 -0.8 2377.2
Kdm6a -0.9 882.8
Olfm2 -2.2 48.7
Dbh -4.3 12.5
Mup10 -5.1 11.9
Xist -9.4 1805.9
log2FC
AveExpr
♂ vs ♀
8-Weeks 4nqo
vs
No Treatment
B
Signature
Stress
Cellular
Metabolic
response to toxic substance
detoxification of inorganic compound
stress response to metal ion
formation of translation preinitiation complex
I-KB phosphorylation
intracellular sodium ion homeostasis
nucleoside bisphosphate metabolic process
ribonucleoside bisphosphate metabolic process
purine nucleoside bisphosphate metabolic process
aromatic AA family catabolic process
-log10(pval.adj)
1.4
1.3
1.2
1.1
1
GeneRatio
0.10
0.14
0.20
8-Weeks 4nqo
vs
No Treatment
♂ vs ♀

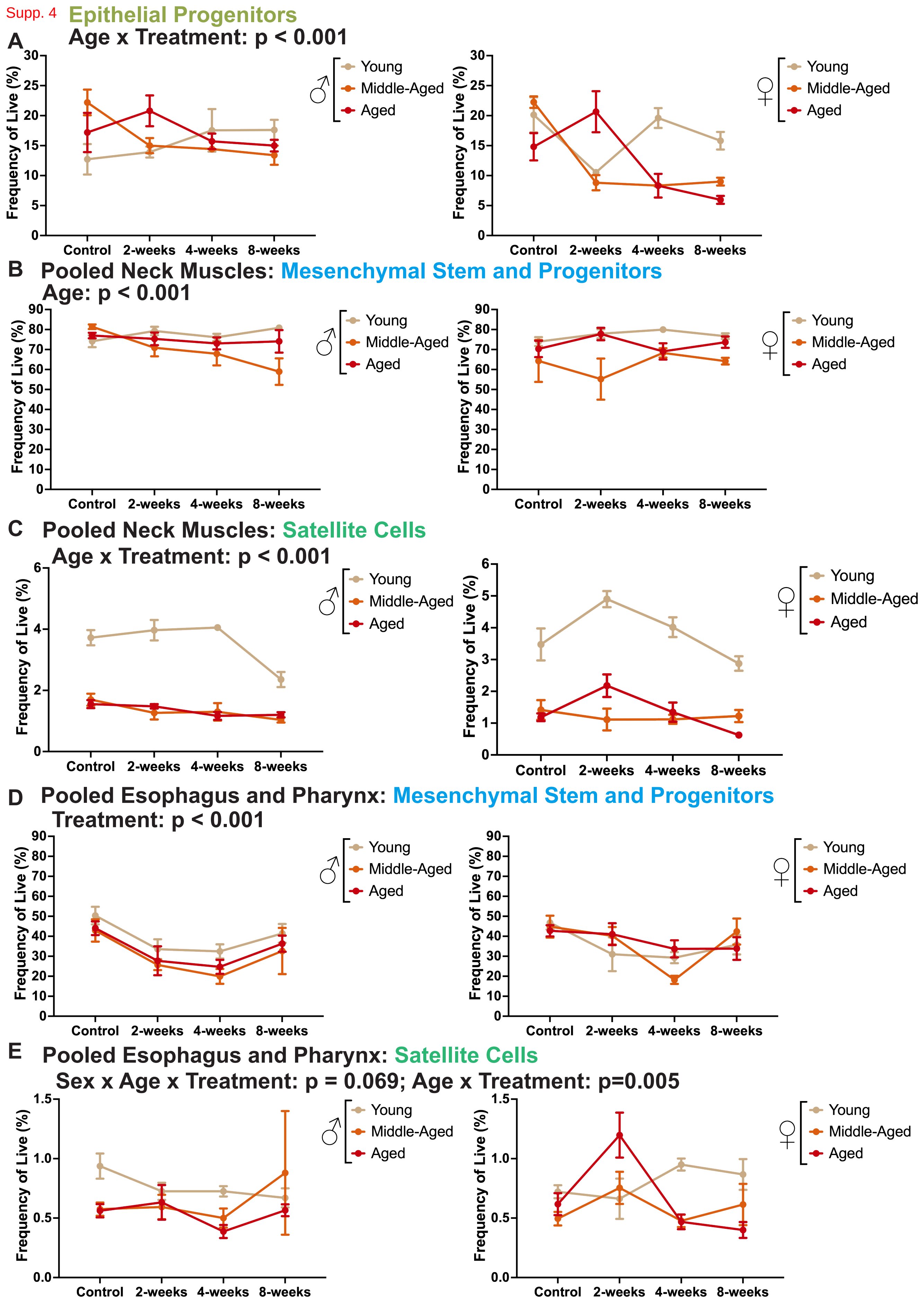
Supp. 4
Epithelial Progenitors
A
Age x Treatment: p < 0.001
Frequency of Live (%)
Control
2-weeks
4-weeks
8-weeks
♂
♀
Young
Middle-Aged
Aged
B
Pooled Neck Muscles: Mesenchymal Stem and Progenitors
Age: p < 0.001
C
Pooled Neck Muscles: Satellite Cells
Age x Treatment: p < 0.001
D
Pooled Esophagus and Pharynx: Mesenchymal Stem and Progenitors
Treatment: p < 0.001
E
Pooled Esophagus and Pharynx: Satellite Cells
Sex x Age x Treatment: p = 0.069; Age x Treatment: p=0.005
